# Isoprenoid biosynthesis regulation in poplars by methylerythritol phosphate and mevalonic acid pathways

**DOI:** 10.1101/2020.07.22.216804

**Authors:** Ali Movahedi, Hui Wei, Boas Pucker, Tingbo Jiang, Weibo Sun, Dawei Li, Liming Yang, Qiang Zhuge

**Author notes:** These authors contributed equally as the first author. (A.M.); (H.W.); (W.S.); (D.L.); (L.Y.); (Q.Z.). (B.P.). (T.J.).

## Abstract

The isoprenoids found in plants are extremely important to survive with various human applications, such as flavoring, fragrance, dye, pharmaceuticals, and biomass used for biofuels. Methylerythritol phosphate (MEP) and mevalonic acid (MVA) pathways are critical in plants, responsible for isoprenoid biosynthesis. 1-deoxy-D-xylulose5-phosphate synthase (DXS) and 1-deoxy-D-xylulose5-phosphate reductoisomerase (DXR) catalyze the rate-limiting steps in the MEP pathway, while 3-hydroxy-3-methylglutaryl-CoA reductase (HMGR) catalyzes the rate-limiting step in the MVA pathway. Here, we showed while *PtHMGR* overexpressors (OEs) exhibited different MEP- and MVA-related gene expressions compared with non-transgenic poplars (NT), the *PtDXR-OEs* revealed upregulated MEP-related and downregulated MVA-related gene expressions. *PtDXR* and *PtHMGR o*verexpressions caused changes in MVA-derived trans-zeatin-riboside, isopentenyl adenosine, castasterone, and 6-deoxocastasterone well as MEP-derived carotenoids and gibberellins. In *PtHMGR*-OEs, the accumulated geranyl diphosphate synthase (*GPS*) and geranyl pyrophosphate synthase (*GPPS)* transcript levels in the MEP pathway led to an accumulation of MEP-derived isoprenoids. In contrast, upregulation of farnesyl diphosphate synthase (*FPS*) expression in the MVA pathway contributed to increased levels of MVA-derived isoprenoids. In addition, *PtHMGR*-OEs increased MEP-related *GPS* and *GPPS* transcript levels, expanded MEP-derived isoprenoid levels, changed *FPS* transcript levels, and affected MVA-derived isoprenoid yields. These results demonstrate the contribution of MVA and MEP pathways regulating isoprenoid biosynthesis in poplars.

## 1. Introduction

Biosynthesis of isoprenoids (terpenoids) is essential for all living organisms. There are over 50,000 distinct molecules in living organisms that are isoprenoids, presenting many functional and structural properties (Thulasiram et al., 2007). Isoprenoids play vital roles in plant growth and development, as well as in membrane fluidity, photosynthesis, and respiration. As specific metabolites, they join in plant-pathogen and allelopathic interactions to preserve plants upon pathogens and herbivores, and they are also created to draw pollinators and seed-dispersing animals. Numerous isoprenoids are of commercial importance for rubber products and drugs, flavors, fragrances, agrochemicals, nutraceuticals, disinfectants, and pigments (Bohlmann and Keeling, 2008). A wide range of isoprenoid biochemical processes are involved in photosynthesis in plants, including electron transfer, quenching of excited chlorophyll triplets, light-harvesting, and energy conversion (Malkin R, 2000). Chlorophylls, consisting of the heme pathway-derived tetrapyrrole ring with an appended isoprenoid-derived phytol chain, exist in all reaction center and antenna complexes to absorb light energy and transfer electrons to the reaction centers. The linear or partially cyclized carotenes and their oxygenated derivative xanthophyll are isoprenoids that extinguish excess excitation energy through light-harvesting to preserve the light-harvesting system. However, these isoprenoids operate as attractants in flowers and fruits as well (Rodriguez-Concepcion, 2010). A large proportion of the isoprenoid flux in plants is conducted toward the synthesis of membrane sterol lipids. In contrast to vertebrates synthesizing cholesterol, higher plants synthesize a complex mix of sterol lipids called phytosterols (Boutte and Grebe, 2009).

Plants isoprenoids include gibberellins (GAs), carotene, Lycopene, cytokinins (CKs), strigolactones (GRs), and brassinosteroids (BRs) are produced through methylerythritol phosphate (MEP) and mevalonic acid (MVA) pathways (Henry et al., 2015; van Schie et al., 2006; Xie et al., 2008). The mentioned pathways are involved in plant growth, development, and response to environmental changes (Bouvier et al., 2005; Kirby and Keasling, 2009). The isopentenyl diphosphate isomerase (IDI) catalyzes the conversion of the isopentenyl diphosphate (IPP) into dimethylallyl diphosphate (DMAPP), leading to provide the basic materials for all isoprenoid productions (Hemmerlin, 2012; Lu et al., 2012; Zhang et al., 2019). The produced IPP and DMAPP play essential roles in MEP and MVA pathways crosstalk (Huchelmann et al., 2014; Liao et al., 2016). The MVA pathway reactions appear in the cytoplasm, endoplasmic reticulum (ER), and peroxisomes (Cowan et al., 1997; Roberts, 2007), producing sesquiterpenoids and sterols. The 3-hydroxy-3-methylglutaryl-CoA reductase (HMGR), a rate-limiting enzyme in the MVA pathway, catalyzes 3-hydroxy-3-methylglutary-CoA (HMG-CoA) to form MVA (Cowan et al., 1997; Roberts, 2007).

Reactions of the MEP pathway occur in the chloroplast and produce carotenoids, GAs, and diterpenoids. 1-deoxy-D-xylulose5-phosphate synthase (DXS) and 1-deoxy-D-xylulose5-phosphate reductoisomerase (DXR) are rate-limiting enzymes in the MEP pathway that catalyze the conversion of D-glyceraldehyde3-phosphate (D-3-P) and pyruvate into 2-C-methyl-D-erythritol4-phosphate (MEP) (Cordoba et al., 2009; Perreca et al., 2020; Wang et al., 2012; Yamaguchi, 2018). Terpenoids like phytoalexin and volatile oils play essential roles in plant growth, development, and disease resistance (Hain et al., 1993; Ren et al., 2008). Photosynthetic pigments convert organic carbon into plant biomass (Esteban et al., 2015). In addition to an extensive range of natural functions in plants, terpenoids also consider the potential for biomedical applications. Paclitaxel is one of the most effective chemotherapy agents for cancer treatment, and artemisinin is an anti-malarial drug (Kim et al., 2016a; Kong and Tan, 2015).

Previous metabolic engineering studies have proposed strategies to improve the production of specific metabolites in plants (Ghirardo et al., 2014; Opitz et al., 2014). For example, PMT and H6H encoding the putrescine N-methyltransferase and hyoscyamine 6 β-hydroxylase respectively produced significantly higher scopolamine in transgenic henbane hairy root. Also, HCHL encoding p-hydroxycinnamoyl-CoA hydratase/lyase accumulated the glucose ester of p-hydroxybenzoic acid (pHBA) in Beta vulgaris hairy root (Rahman et al., 2009; Zhang et al., 2004). The 3-hydroxy-3-methylglutaryl-coenzyme A synthase (HMGS) is the second enzyme in the MVA pathway. Liao et al. (2018) confirmed that *HMGS* overexpression of *Brassica juncea* upregulates carotenoid and phytosterol in tomatoes. HMGR has been considered a critical factor in metabolically engineering terpenoids (Aharoni et al., 2005; Dueber et al., 2009). In addition, *PgHMGR1* overexpression of ginseng increases ginsenosides content, a necessary pharmaceutically active component (Kim, 2014).

Transgenic tobacco overexpressing the *Hevea brasiliensis HMGR* enhanced the phytosterol levels (Schaller et al., 1995). It has been shown (Dai et al., 2011) that *SmHMGR2* in *Salvia miltiorrhiza*, resulting in the improvement of squalene and tanshinone contents. Moreover, *Arabidopsis thaliana HMGR1 (AtHMGR1)* enhanced the phytosterol levels in the first generation of transgenic tomatoes (Enfissi et al., 2005). While the deaccumulation of *DXR* transcripts resulted in lower pigmentation and chloroplast appearance defects, the upregulated *DXR* expression caused the MEP-derived plastid isoprenoids to accumulate. Therefore, *DXR* can be genetically engineered to regulate the content of terpenoids and expressed *DXR* in *Arabidopsis* and observed enhanced flux through the MEP pathway (Carretero-Paulet et al., 2006). While the *A*. *thaliana DXR* overexpression caused the diterpene anthiolimine to accumulate in *Salvia sclarea* hairy roots (Vaccaro et al., 2014), the peppermint *DXR* overexpression resulted in essential oil inflation (about 50%) with no significant variations in monoterpene composition (Mahmoud and Croteau, 2001). Furthermore, previous studies have shown the exchange of metabolic intermediates included in the MVA- and MEP pathways through plastid membranes (Laule, 2003; Liao, 2006). In summary, the overexpression of genes involved in the MVA- and MEP pathways can change the abundances or activities of related enzymes and metabolic products, causing a new opportunity for plant breeding to enhance the accumulation of related metabolic products.

Poplars as an economic and energy species are widely used in industrial and agricultural production. Its fast growth characteristics and advanced resources in artificial afforestation play a vital role in the global ecosystem (Devappa, 2015).

This study investigates the poplar isoprenoid biosynthesis. We showed that the overexpression of the MVA-specific *PtHMGR* gene upregulated not only MVA- but MEP-related genes in the transcript levels. We also proved that the overexpression of the MEP-specific *PtDXR* gene caused downregulating MVA-related genes compared with upregulating MEP-related genes, enhancing terpenoid accumulation. Taken together, these results indicate that the MEP is a dominant pathway in contribution with the MVA pathway to produce isoprenoids secondary metabolites, and *HMGR* and *DXR* genes play key regulation points in these pathways.

## 2. Materials and Methods

### 2.1. Plant materials and growth conditions

Non-transgenic *P*. *trichocarpa* and *Populus* × *euramericana* cv. ‘Nanlin 895′ plants were cultured in half-strength Murashige and Skoog (1/2 MS) medium (pH 5.8) under conditions of 24°C and 74% humidity (Movahedi et al., 2015). Subsequently, NT and transgenic poplars were cultured in 1/2 MS under long-day conditions (16 h light/8 h dark) at 24°C for 1 month (Movahedi et al., 2018).

### 2.2. PtHMGR and PtDXR genes isolation and vector construction

To produce cDNA, total RNA was extracted from *P. trichocarpa* leaves and processed with PrimeScript™ RT Master Mix, a kind of reverse transcriptase (TaKaRa, Japan). Forward and reverse primers (Supplementary Table 1: *PtHMGR*-F and *PtHMGR*-R) were designed, and the open reading frame (ORF) of *PtHMGR* was amplified via PCR. We then used the total volume of 50μl including 2 μl primers, 2.0 μl cDNA, 5.0 μl 10 × PCR buffer (Mg2+), 4μl dNTPs (2.5 mM), 0.5 μl rTaq polymerase (TaKaRa, Japan) for the following PCR reactions: 95°C for 7 min, 35 cycles of 95°C for 1 min, 58°C for 1 min, 72°C for 1.5 min, and 72°C for 10 min. Subsequently, the product of the *PtHMGR* gene was ligated into the pEASY-T3 vector (TransGen Biotech, China) based on blue-white spot screening, and the *PtHMGR* gene was inserted into the vector pGWB9 (Song et al., 2016) using Gateway technology (Invitrogen, USA). On the other hand, all steps to generate cDNA, RNA extraction, PCR, pEASY-T3 ligation, and vector construction (pGWB9-PtDXR) of *PtDXR* have been carried out according to Xu et al. (2019).

### 2.3. phylogenetic analyses

We applied the ClustalX for multiple sequence alignment of HMGR proteins, and MEGA5.0 software was used to construct a phylogenetic tree using 1000 bootstrap replicates. The amino acid sequences of HMGR from *Populus trichocarpa*, *Arabidopsis thaliana*, *Gossypium raimondii*, *Malus domestica*, *Manihot esculenta*, *Oryza sativa*, *Prunus persica*, *Theobroma cacao*, and *Zea mays* were obtained from the National Center for Biotechnology Information database (https://www.ncbi.nlm.nih.gov/) and Phytozome (https://phytozome-next.jgi.doe.gov/).

### 2.4. Transgenic poplars: generation and confirmation

*Agrobacterium tumefaciens* var. EHA105 was used for the infection of poplar leaves and petioles (Movahedi et al., 2014). Poplar buds were screened on differentiation MS medium supplemented with 30 μg/mL Kanamycin (Kan). Resistant buds were planted in bud elongation MS medium containing 20 μg/mL Kan and transplanted into 1/2 MS medium including 10 μg/mL Kan to generate resistant poplar trees. Genomic DNA has been extracted from putative transformants one-month-old leaves grown on a kanamycin-containing medium using TianGen kits (TianGen BioTech, China). The quality of the extracted genomic DNA (250–350 ng/μl) was determined by a BioDrop spectrophotometer (UK). PCR was carried out using designed primers (Supplementary Table 1: CaMV35S as the forward and PtHMGR as the reverse), Easy Taq polymerase (TransGene Biotech), and 50 ng of extracted genomic DNA as a template to amplify about 2000 bp. In addition, total RNA was extracted from these one-month-old leaves to produce cDNA, as mentioned above. These cDNA then were applied to reverse transcription-quantitative PCR (RT-qPCR) (Supplementary Table 1: PtHMGR forward and reverse) for comparing the transformants *PtHMGR-OEs* expressions with NT poplars and transforming confirmation.

### 2.5. Phenotypic properties evaluation

To evaluate phenotypic changes, we selected 45-day-old poplars from PtHMGR-and PtDXR-OEs and NT poplars. We then simultaneously calculated the stem lengths (mm) and stem diameters (mm) every day and recorded them. All recorded were analyzed by GraphPad Prism 9, applying ANOVA one way (Supplementary Table 2).

### 2.6. Analyses via qRT-PCR

12-month-old *PtDXR-OEs* (Xu et al., 2019) and *PtHMGR-OE* poplars (Soil-grown poplars) have been used to extract total RNA. The qRT-PCR was performed to identify MVA- and MEP-related gene expression levels in NT, *PtDXR-OE*, and *PtHMGR-OE* poplars. The qRT-PCR was served with a StepOne Plus Real-time PCR System (Applied Biosystems, USA) and SYBR Green Master Mix (Roche, Germany). Poplar *Actin* (*PtActin*) (XM-006370951.1) was previously tested as a reference gene for this experiment (Zhang et al., 2013). The following conditions were used for qRT-PCR reactions: pre-denaturation at 95°C for 10 min, 40 cycles of denaturation at 95°C for 15 s, and a chain extension at 60°C for 1 min. Three independent experiments were conducted using gene-specific primers (Supplementary Table 1: PtHMGR forward and reverse).

### 2.7. Metabolite analyses via high-performance liquid chromatography-tandem mass spectrometry

The isopropanol/acetic acid extraction method extracted poplar endogenous hormones from NT, *PtDXR-OE*, and *PtHMGR-OE* leaves. GAs and CKs were extracted from, and then HPLC-MS/MS (Qtrap6500, Agilent, USA) was used to quantify levels of GAs, zeatin, tZR, and IPA. Also, methanol considered as solvent was used to extract 5-Deoxystrigol (5-DS), CS, and DCS, and HPLC-MS/MS (Aglient1290, AB; SCIEX-6500Qtrap, Agilent; USA) was also used to determine the contents of 5-DS, CS, and DCS. In addition, acetone, as a solvent, was used to isolate the carotenoid component of poplar leaves. To identify the carotenoid contents, the peak areas of carotenoids analyzed by HPLC (Symmetry Shield RP18, Waters, USA) were used to draw standard carotenoid curves, including β-carotene, Lycopene, and Lutein. Also, the HPLC was used to determine the contents of carotenoids, including β-carotene, Lycopene, and Lutein in NT and OE lines.

## 3. Results

### 3.1. Identification, analyses, and Isolation of PtHMGR and PtDXR genes

*Populus trichocarpa* v3.1 (Phytozome genome ID: 444, NCBI taxonomy ID: 3694) has been applied to download 595 amino acids (aa) PtHMGR (Potri.004G208500.1) and the other species’ HMGR to align and analyze. High similarity, including lots of conserved aa accompanied by specific similar domains, HMG-CoA-binding motifs (EMPVGYVQIP’ and ‘TTEGCLVA), and NADPH-binding motifs (DAMGMNMV’ and ‘VGTVGGGT) (Ma et al., 2012) (Supplementary Figure 1), confirmed the PtHMGR protein analytically. Consequently, a phylogenetic tree based on the various species HMGR supported the PtHMGR candidate identification (Supplementary Figure 2). The tblastn was then applied to reveal 2614 bp *PtHMGR* located on Chr04:21681480..21684242 with a 1785 bp CDS. After that, the amplified 1857 bp of the *PtHMGR* from *Populus trichocarpa* cDNA confirmed the putative transgenic lines (Supplementary Figure 3a), exhibiting amplicons in PCR identification compared to NT poplar (Supplementary Figure 3b). The transgenic poplars (*PtHMGR-OEs*) also showed enhanced *PtHMGR* expressions than NT (Supplementary Figure 3c), indicating successful overexpression of *PtHMGR* in poplar. The *PtDXR* gene, which has been isolated, sequenced, and analyzed previously by the authors (Xu et al., 2019), was then transferred into poplar genome to generate PtDXR-OEs used in this study.

### 3.2. PtHMGR- and PtDXR overexpressions regulate MVA-related gene expressions

MVA-related genes *AACT*, *MVK*, *MVD, and FPS*, except *HMGS*, were significantly upregulated in *PtHMGR-OE* transgenics than NT poplars (Supplementary Figure 4a). In contrast, while only *FPS* revealed significant upregulation by *PtDXR-OEs* in transgenics compared with NT, the other MVA-related genes *AACT*, *HMGS*, *HMGR*, and *MVK* were considerably downregulated (Supplementary Figure 4a). However, the mean comparison of MVA-related gene expressions regulated by *HMGR* and *DXR* overexpressing exhibited significant upregulated *MVK* through *PtHMGR-OEs* (Figure 1a). The *HMGS* was significantly downregulated within *PtHMGR*-and *PtDXR-OEs*, and *MVK* was downregulated by *PtDXR* overexpression (Figure 1a). The mean comparison of *AACT*, *MVD*, and *FPS* revealed upregulation through *PtHMGR-OEs*, while *AACT* and *MVD* showed downregulation through *PtDXR-OEs* (Figure 1a).

**Figure 1.**
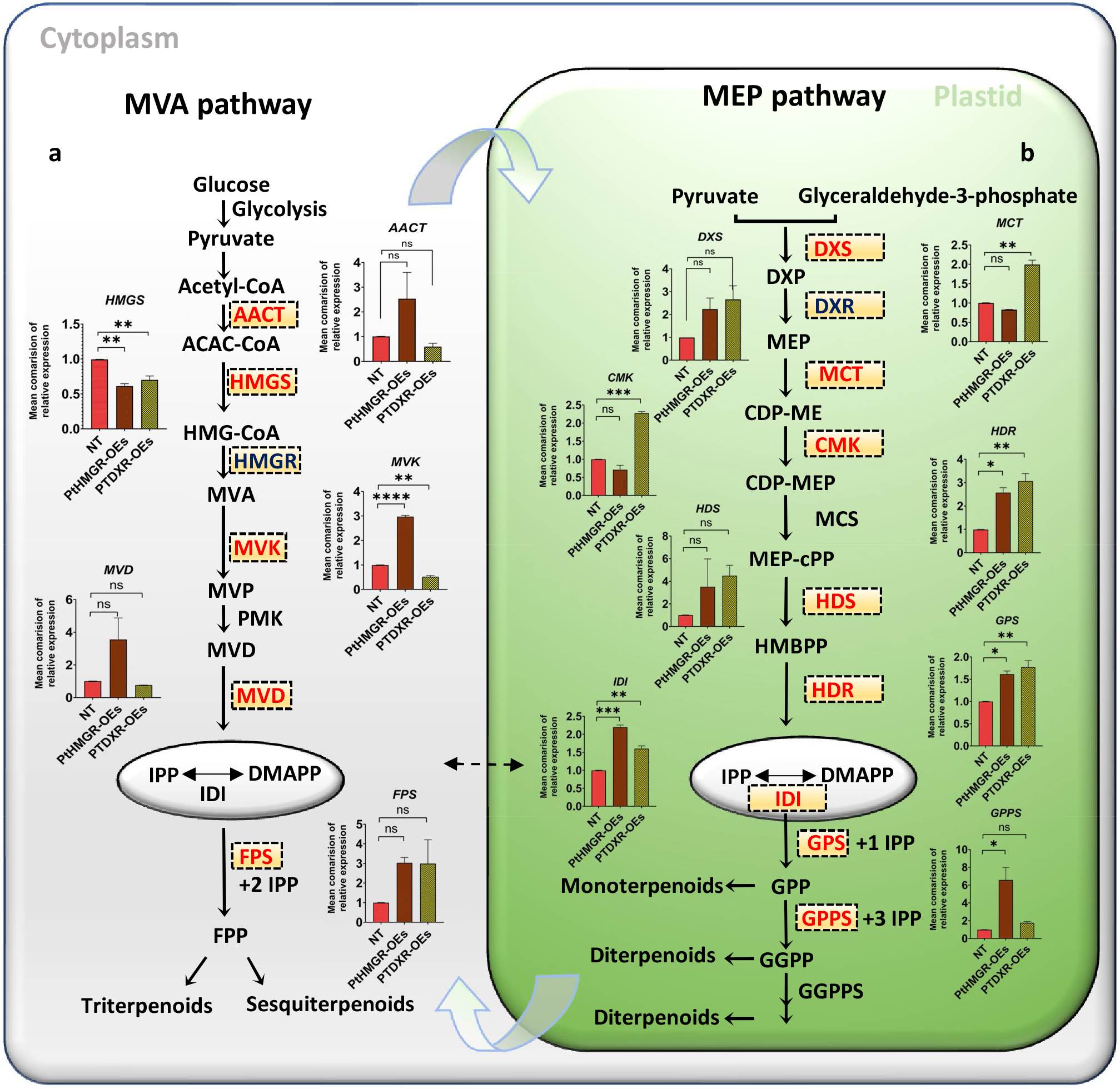
MVA- and MEP-related genes analyses in overexpressed *PtHMGR*- and *PtDXR-OEs* poplars. **a**, Mean comparison of relative expression of MVA-relate genes *AACT*, *HMGS*, *MVK*, *MVD*, and *FPS* (Indicated in red) affected by *PtHMGR* overexpressing. **b**, Mean comparison of relative expression of MEP-related genes *DXS*, *MCT*, *CMK*, *HDS*, *HDR*, *IDI, GPS*, and *GPPS* (Indicated in red) affected by *PtHMGR* overexpressing; *HMGR* and *DXR*, which were overexpressed respectively by *DXR*- and *HMGR-OEs*, were presented in Supplementary Figure 4. *PtActin* was used as an internal reference in all repeats; “ns” means not significant, * P < 0.05, ** P < 0.01, ***P < 0.001, ****P < 0.0001; Three independent replications were performed in this experiment.

### 3.3. PtHMGR- and PtDXR overexpressions regulate MEP-related gene expressions

While, the expression of MEP-related genes *DXS*, *DXR*, 1-hydroxy-2-methyl-2-(E)-butenyl-4-diphosphate synthase (*HDS*), 1-hydroxy-2-methyl-2-(E)-butenyl-4-diphosphate reductase (*HDR*), *IDI*, and *GPPS* were significantly upregulated in all *PtHMGR-OEs* transgenic poplars in comparison with NT, the *GPS* overexpression was enhanced only by *PtHMGR-OE3* (Supplementary Figure 4c). In addition, 2-C-methyl-d-erythritol4-phosphate cytidylyltransferase (*MCT*) and 4-diphosphocytidyl-2-C-methyl-D-erythritol kinase (*CMK*) have been downregulated by *PtHMGR-OEs* (Supplementary Figure 4c). In contrast, all MEP-related genes were upregulated significantly by *PtDXR* overexpression (Supplementary Figure 4d). In total, the comparison of the effect of *PtHMGR* on MEP-related genes exhibited significant upregulations of *HDR*, *IDI*, *GPS*, and *GPPS* (Figure 1b). These comparisons also revealed the upregulation of *DXS* and *HDS* except *MCT* and *CMK*, downregulated by *PtHMGR* overexpression (Figure 1b). In contrast, the mean comparison of the effect of *PtDXR* overexpression on MEP-related genes exhibited upregulation of the all mentioned above genes (Figure 1b).

### 3.4. MVA- and MEP-derived carotenoids are affected by PtHMGR-and PtDXR-OEs

β-carotene is a carotenoid synthesis that has been broadly used in the industrial composition of pharmaceuticals and as food colorants, animal supplies additives, and nutraceuticals. MVA-and MEP pathways have been proved effective in the biosynthesis of β-carotene (Yang, 2014). In addition, Lycopene is a carotenoid referring to C40 terpenoids and is broadly found in various plants, particularly vegetables and fruits. It has been shown that MVA and MEP-pathways directly influence the biosynthesis production of Lycopene (Kim et al., 2019; Wei et al., 2018). While Wille et al. (2004) showed that β-carotene and Lutein are synthesized using intermediates from the MEP pathway, Opitz et al. (2014) revealed that both MVA and MPE pathways contribute to producing isoprenoids such as β-carotene and Lutein. HPLC-MS/MS has analyzed the quantity of MVA and MEP derivatives. Our analyses revealed that *HMGR*-OEs caused a significant enhancement in Lycopene (an average of ~ 0.08 ug/g), β-carotene (an average of ~ 0.33 ug/g), and Lutein (an average of ~ 272 ug/g) production compared with NT poplars (~0.02, ~0.08, and ~100 ug/g respectively) (Figure 2a, b, and c; Supplementary Figure 5). The ABA-related gene expressions also have been calculated. Results revealed a significantly increased *ZEP1*, *2*, and *3* relative gene expressions with averages of ~2.85, ~4.67, and ~2.92 compared to NT with an average of ~1 (Figure 2d). These results also showed meaningful enhancements of *NCED1, 2*, and *3* relative gene expressions with the averages of ~4.16, ~3.79, and ~3.4 compared to NT with an average of ~1 (Figure 2e).

**Figure 2.**
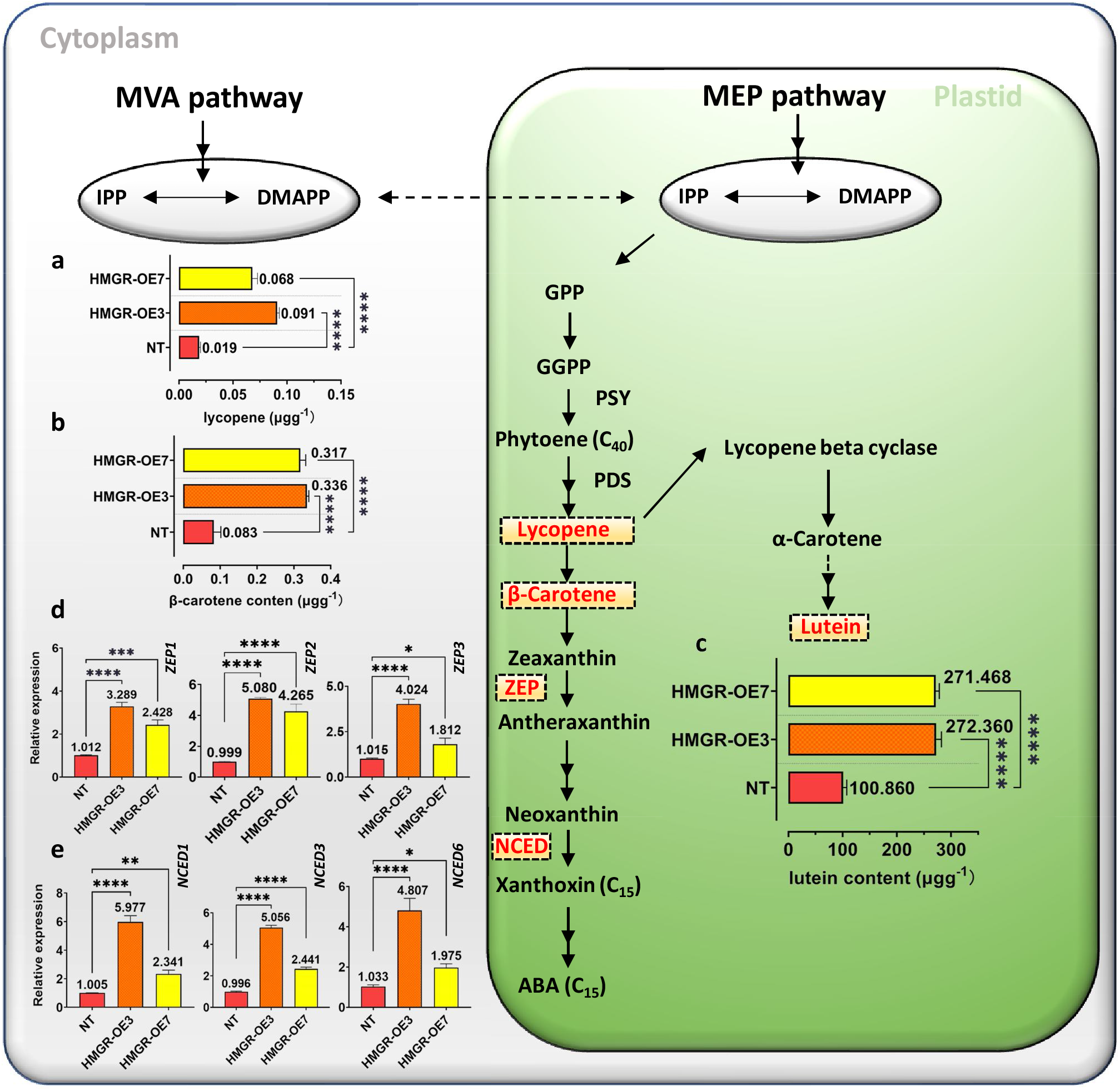
HPLC-MS/MS content analyses of lycopene, β-carotene, lutein, and real-time PCR of *ZEP* and *NCED* genes family. HPLC-MS/MS content analyses have been performed to show the effect of *PtHMGR-OEs* on **a**, lycopene **b**, β-carotene, and **c**, lutein. Relative expressions have been analyzed affected by *PtHMGR-OEs* compared with NT poplars of **d**, *ZEP*, and **e**, *NCED* genes family. Bars represent mean ± SD (n = 3); Stars reveal significant differences, * P < 0.05, ** P < 0.01, *** P < 0.001, ****P < 0.0001; Three independent experiments were performed in these analyses.

On the other hand, the levels of the MEP-derived substances lycopene, β-carotene, and Lutein were significantly increased in *PtDXR*-OEs with the averages of ~0.08, 0.22, 209.32 ug/g, respectively compared to NT poplars (Figure 3a, b, and c; Supplementary Figure 6). The analyses of ABA-related gene expressions revealed significantly increased *ZEP1*, *2*, and *3* relative gene expressions with the averages of ~2.63, ~2.38, and ~3.86 compared to NT with an average of ~1 (Figure 3d). These results also showed meaningful enhancements of *NCED2* and *3* relative gene expressions with averages of ~2.25 and ~2.21 compared to NT with an average of ~1 (Figure 2e). These results revealed a decreased average in *NCED1* relative gene expression with an average of ~0.66 compared to NT poplars.

**Figure 3.**
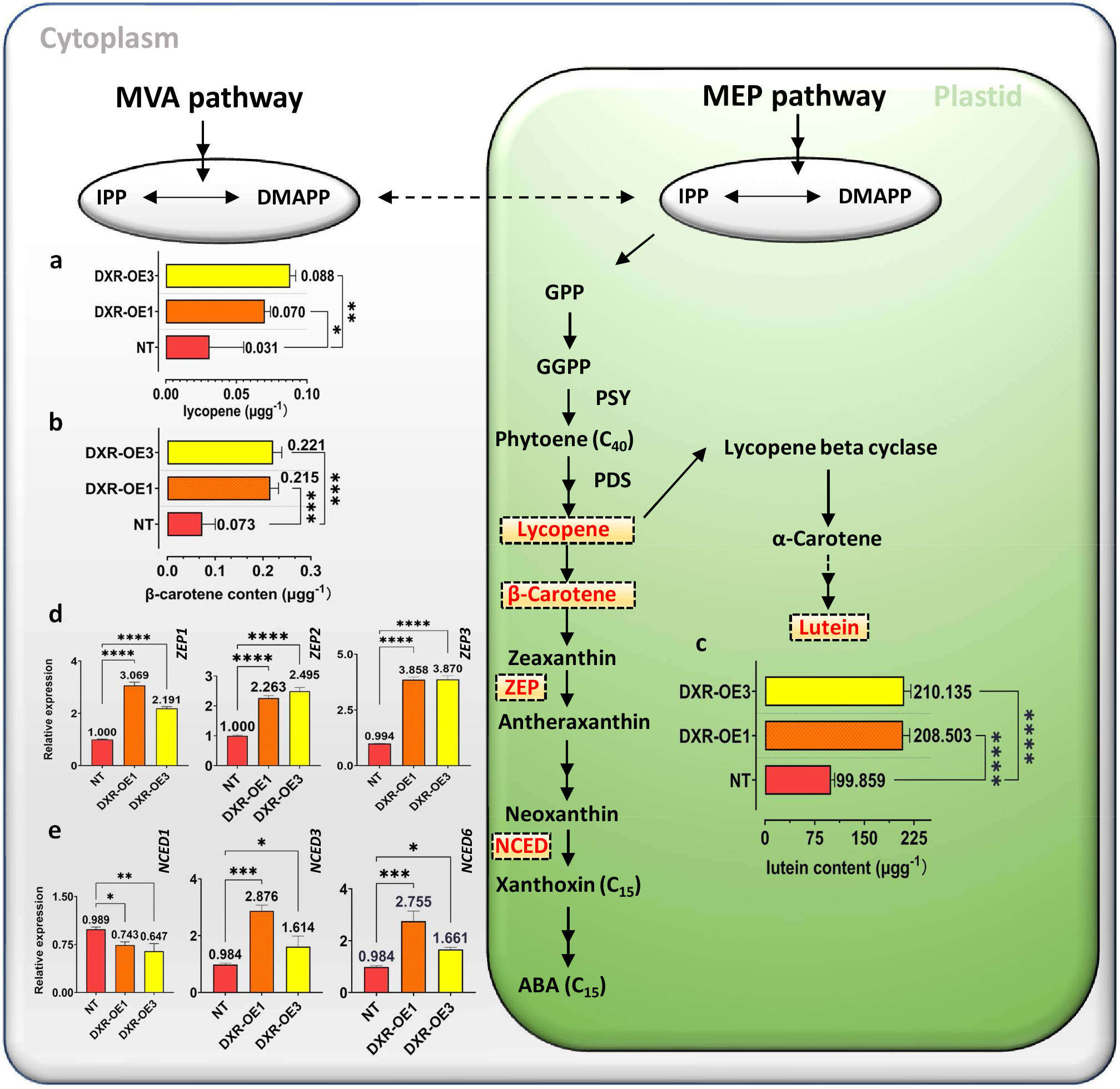
HPLC-MS/MS content analyses of lycopene, β-carotene, lutein, and real-time PCR of *ZEP* and *NCED* genes family. HPLC-MS/MS content analyses have been performed to show the effect of *PtDXR-OEs* on **a**, lycopene **b**, β-carotene, and **c**, lutein. Relative expressions have been analyzed affected by *PtDXR-OEs* compared with NT poplars of **d**, *ZEP*, and **e**, *NCED* genes family. Bars represent mean ± SD (n = 3); Stars reveal significant differences, * P < 0.05, ** P < 0.01, *** P < 0.001, ****P < 0.0001; Three independent experiments were performed in these analyses.

### 3.5. MVA and MEP-related derivatives are influenced by PtHMGR- and PtDXR-OEs

The other MVA and MEP derivatives such as GAs, trans-zeatin-riboside (tZR), isopentenyl adenosine (IPA), 6-deoxyocastasterone (DCS), and castasterone (CS) productions affected by *PtHMGR*- and *PtDXR-OEs* have been analyzed. While Gibberellic acid (GA3) (a downstream product of MEP) ( an average of ~0.22 ng/g), tZR (an average of ~0.06 ng/g), IPA (an average of ~0.59 ng/g), DCS (an average of 4.95 ng/g) revealed significantly more productions induced by *HMGR*-OEs, the CS production (~0.095 ng/g) was decreased considerably compared to NT poplars (~0.10, ~0.03, ~0.37, ~1.50, and ~0.20 ng/g respectively) (Figure 4a–j). These results demonstrate that the *HMGR* gene interacts with MVA and MEP derivatives productions in plants. On the other hand, the *PtDXR* overexpression significantly affected the contents of MEP- and MVA-derived products except for CS. *PtDXR*-OEs showed a significant increase ~0.276 ng/g in the GA3 content (Figure 4a and f). The tZR content represented a 10-fold increase (~0.304 ng/g) affected by *PtDXR-OEs* compared to NT poplars (0.032 ng/g) (Figure 4b and g). The content of IPA in *PtDXR-OEs* meaningfully increased ~ 0.928 ng/g, compared to 0.363 ng/g in NT poplars (Figure 4c and h) with a 3-fold increase. In addition, the DCS content considerably increased to ~3.36 ng/g, compared with ~1.50 ng/g in NT, representing a 3-fold increase in *PtDXR-OEs* (Figure 4d and i). By contrast, the content of CS in *PtDXR-OEs* significantly decreased (~0.137 ng/g) compared to NT poplar (0.203 ng/g), indicating significant down-regulation in *PtDXR-OEs* (Figure 4e and j). The HPLC-MS/MS chromatograms of GA, tZR, IPA, DCS, and CS standards are provided in Supplementary Figures 7–11.

**Figure 4.**
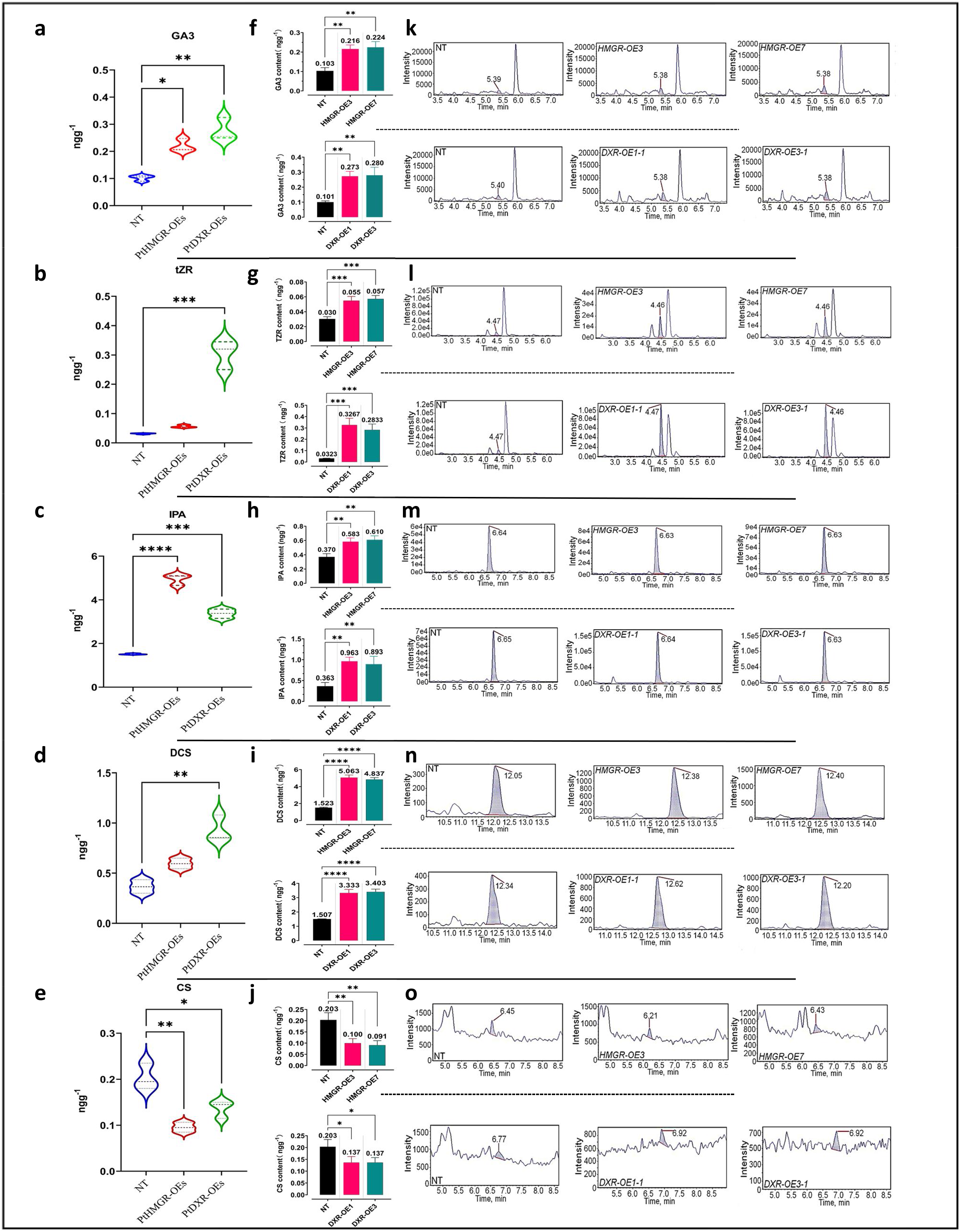
HPLC-MS/MS content analyses of MEP- and MVA-derived isoprenoids. **a,b,c,d**, and **e**, Violin plots reveal the contents of isoprenoids GA3, tZR, IPA, DCS, and CS obtained from MEP- and MVA pathways influenced by *PtHMGR*- and *PtDXR-OEs*. **f,g,h,i, and j,** the column plots reveal the effect of *PtHMGR-OE3* and −*7* and *PtDXR-OE1* and −*3* on the mentioned above isoprenoids separately; NT poplars have been used as the control. Bars represent mean ± SD (n = 3); Stars reveal significant differences, *P < 0.05, **P < 0.01, ***P < 0.001, ****P < 0.0001. **k,l,m,n, and o,** represent the HPLC-MS/MS chromatogram content analyses of GA3, tZR, IPA, DCS, and CS, respectively affected by *PtHMGR- and PtDXR-OEs* compared with NT poplars.

### 3.6. Phenotypic properties

To investigate the growth and development resulting from different produced isoprenoids contents amongst the affected MVA-and MEP pathways contributions by *PtHMGR*-and *PtDXR-OEs*, we decided to evaluate phenotypic stem lengths and diameters changes. Results exhibited a significant increase in GA3 contents in *PtDXR-OEs* (Figure 4a) associated with a considerable rise in cytokinin tZR (Figure 4b), resulting in significantly more development in stem length compared to *PtHMGR-OEs* and NT poplars (Figure 5a and b). Regarding increasing ABA-related gene expressions (*ZEP* and *NCED*) in *PtHMGR-OEs* than *PtDXR-OEs* and NT poplars (Figure 5c and d) and also concerning insufficient increase cytokinin tZR in *PtHMGR-OEs* compared with NT poplars (Figure 4b), *PtHMGR* transgenics showed a shorter stem length that *PtDXR* transgenics compared with NT poplars (Figure 5a and b). We also observed that only *PtDXR-OEs* revealed a few significant increases in stem diameters than PtHMGR-OEs and NT poplars (Figure 5e).

**Figure 5.**
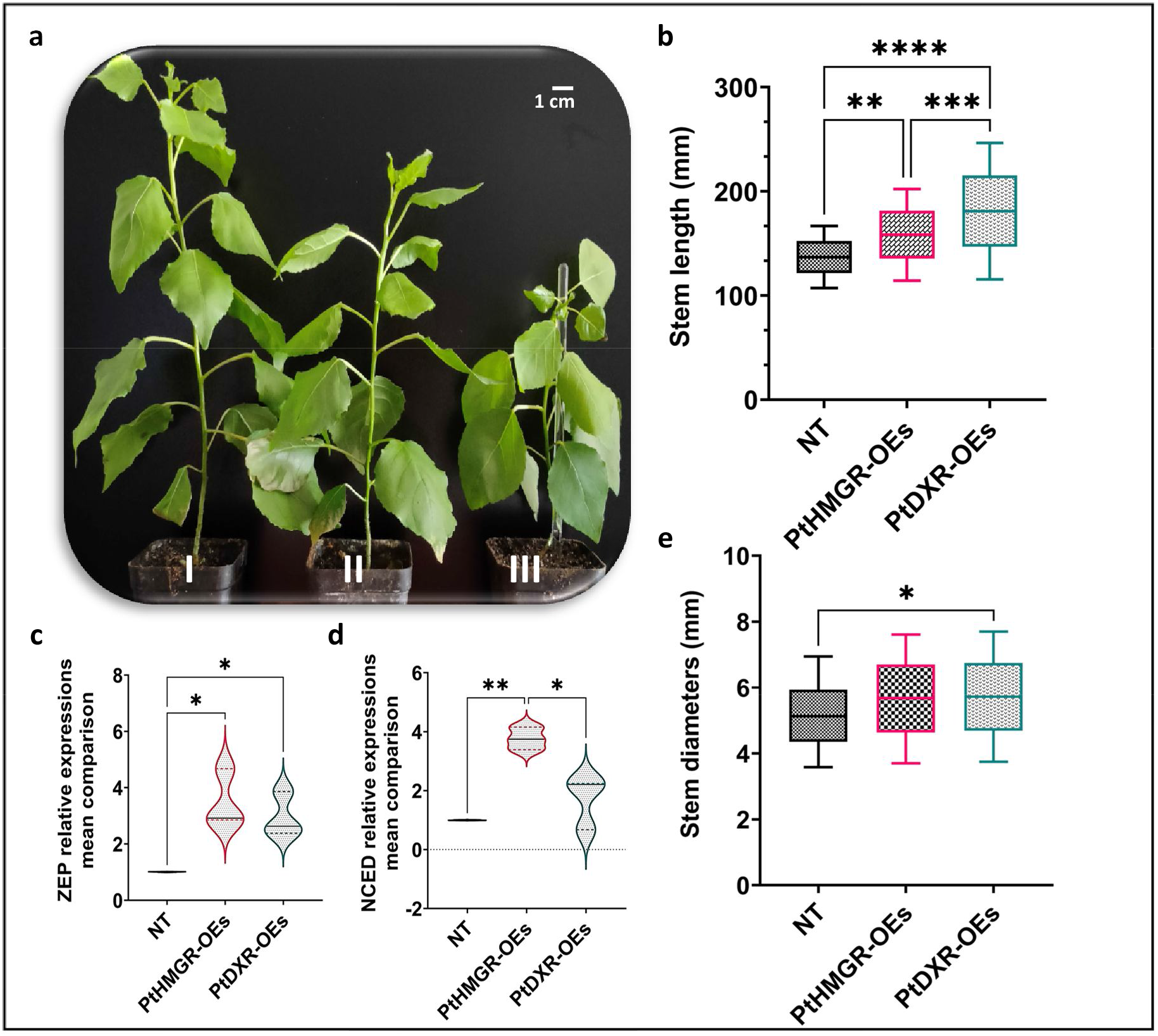
Phenotypic changes resulted by affected communications of MVA- and MEP pathways amongst *PtHMGR*- and *PtDXR-OEs* in 45-day-old poplars. **a**(I), The PtDXR transgenic revealed a higher stem length than *PtHMGR-OEs* and NT poplars. **a**(II), The *PtHMGR* transgenic presents an insignificantly more stem development than NT poplar. **a**(III), NT poplar was used as a control; Scale bar represents 1 cm. **b**, The Box and Whisker mean comparison plot of stem lengths revealed significantly higher lengths *PtDXR-OEs* than NT poplars compared with *PtHMGR-OEs*. *PtHMGR* transgenics also revealed significantly higher lengths than NT poplars. **c** and **d**, The Violin mean comparison plots of *ZEP* and *NCED* relative expressions between *PtHMGR*-and *PtDXR-OEs* compared to NT poplars. **e**, The Box and Whisker mean comparison plot of stem diameters revealed less significant differences between *PtDXR-OEs* and NT poplars. Stars reveal significant differences, *P < 0.05, **P < 0.01, ***P < 0.001, ****P < 0.0001.

## 4. Discussion

### 4.1. The HMGR and DXR crucial roles in isoprenoid biosynthesis

Several studies report that HMGR activity is regulated by isoprenoid outcomes when stigmasterol and cholesterol reduce the HMGR activity by 35% (Russell and Davidson, 1982). Utilization of the isoprenoid growth control abscisic acid also prevented HMGR activity in pea by about 40%, while zeatin and gibberellin, other isoprenoid growth regulators, improved the activity of HMGR (Russell and Davidson, 1982). Maurey et al. (1986) reported that the alga *Ochromonas malhamensis* developed in mevinolin exhibited to 15-fold increase in microsomal HMGR activity, slightly influencing cell growth. Moreover, the MEP pathway is the primary precursor for required plastid isoprenoids (Wright et al., 2014). It has been shown that volatile compounds made by the MEP pathway are involved in plant protection against biotic and abiotic stresses (Gershenzon and Dudareva, 2007). By modifying the expression of DXR, promising metabolite compounds have developed in the mint plant (Mahmoud and Croteau, 2002). In addition, the DXR overexpression in Arabidopsis resulted in accumulating isoprenoids such as tocopherols, carotenoids, and chlorophylls (Carretero-Paulet et al., 2006). DXR overexpression has also been proven to improve diterpene contents in transgenic bacteria (Morrone et al., 2010). Biotic stresses are vital in providing pharmaceutical terpenoids by expanding the number of enzymes included in biosynthetic pathways by controlling biosynthetic genes expression (Kang et al., 2009; Lu et al., 2016). Biotic stresses caused to improve DXR expression followed by triptophenolide content in *Tripterygium wilfordii* cell culture suspension (Tong et al., 2015).

### 4.2. Overexpression of PtDXR results in upregulation of isoprenoid biosynthesis gene expression

Liao et al. (2018) showed that overexpression of *BjHMGS1* affects the expression levels of MEP- and MVA-related genes and slightly increases the transcript levels of *DXS* and *DXR* in transgenic plants. However, *DXS*, *DXR*, *HDS*, and *HDR* expression levels have been upregulated significantly in *PtHMGR-OE* poplars, while *MCT* and *CMK* are downregulated.

Similar to Liao et al. (2018) which the *BjHMGS1* overexpression in tomatoes significantly increased the *GPS* and *GPPS* expressions, we exhibited that the *PtHMGR* overexpression enhanced the farnesyl diphosphate synthase (*FPS*), *GPS*, and *GPPS* expressions may stimulate the crosstalk between IPP and DMAPP, increasing the biosynthesis of plastidial C15 and C20 isoprenoid precursors. Xu et al. (2012) showed that *HMGR* overexpression in *Ganoderma lucidum* caused upregulated *FPS*, squalene synthase (*SQS*), or lanosterol synthase (LS) mRNA expressions and developed the contents of ganoderic acid and intermediates, including squalene and lanosterol. In addition, the *BjHMGS1* overexpression in tomatoes significantly increased transcript levels of *FPS*, *SQS*, squalene epoxidase (*SQE*), and cycloartenol synthase (*CAS*) (Liao et al., 2018). This study exhibited that except for *HMGS* downregulating, the *AACT*, *MVK*, and *MVD* transcript levels were significantly upregulated in *PtHMGR-OE* poplars. We revealed that these enhanced gene expressions mainly were associated with the MVA-related genes contributing to the biosynthesis of sesquiterpenes and other C15 and universal C20 isoprenoid precursors.

### 4.3. Overexpression of PtDXR affects MEP- and MVA-related genes

Zhang et al. (2018) showed that the *TwDXR* overexpression in *Tripterygium wilfordii* increases the *TwHMGS*, *TwHMGR*, *TwFPS*, and *TwGPPS* expressions but decreases the *TwDXS* expression. Moreover, Zhang et al. (2015) exhibited that the *NtDXR1* overexpression in tobacco increases the transcript levels of eight MEP-related genes, indicating that the *NtDXR1* overexpression led to upregulated MEP-related gene expressions. In *A. thaliana*, the *DXR* transcript level changes do not affect *DXS* gene expression or enzyme accumulation, although the *DXR* overexpression promotes MEP-derived isoprenoids such as carotenoids, chlorophylls, and taxadiene (Carretero-Paulet et al., 2006).

On the other hand, the potato *DXS* overexpression in *A. thaliana* led to upregulation of downstream *GGPPS* and phytoene synthase (*PSY*) genes (Henriquez et al., 2016). Furthermore, (Simpson et al., 2016) exhibited that the *A. thaliana DXS* overexpression in Daucus carota caused to enhance the *PSY* expression significantly.

In this study, while the *PtDXR-OEs* exposed higher MEP-related gene expressions than NT poplars, the *PtDXR-OEs* revealed significant downregulated MVA-related gene expressions than NT poplars. These findings illustrate that the MEP pathway regulates monoterpenes, diterpenes, and tetraterpenoids biosynthesis and could affect the MVA pathway.

The diversity of biosynthetic pathways, the complexity of metabolic networks, and the insufficient knowledge of gene regulation led to species-specific regulation patterns of MEP- and MVA-related gene expression. One possible conclusion is that MEP- and MVA-related genes often do not work alone but are co-expressed with upstream and downstream genes in the MEP- and MVA- pathways to carry out a specific function.

### 4.4. Overexpression of HMGR promotes the formation of GAs, and carotenoids in plastids and accumulation of tZR, IPA, and DCS in the cytoplasm

HMGR, as the rate-limiting enzyme in the MVA-pathway of plants, plays a critical role in controlling the flow of carbon within this metabolic pathway. The upregulation of *HMGR* significantly increases isoprenoid levels in plants. Overexpression of *HMGRs* of different plant species has been reported to raise isoprenoids levels significantly. The heterologous expression of *Hevea brasiliensis HMGR1* in tobacco increased the sterol content and accumulated intermediate metabolites (Schaller et al., 1995). The *A*. *thaliana HMGR o*verexpression in *Lavandula latifolia* increased the levels of sterols in the MVA-and MEP-derived monoterpenes and sesquiterpenes (Munoz-Bertomeu et al., 2007). In addition, the *Salvia miltiorrhiza SmHMGR* overexpression in hairy roots developed MEP-derived diterpene tanshinone (Kai et al., 2011). In our study, ABA synthesis-related genes (*NCED1*, *NCED3*, *NCED6*, *ZEP1*, *ZEP2*, and *ZEP3*) and the contents of GA3 and carotenoids were upregulated in *PtHMGR-OE* poplar seedlings. This finding suggests that the *HMGR* overexpression may indirectly affect the biosynthesis of MEP-related isoprenoids, including GA3 and carotenoids. The accumulation of MVA-derived isoprenoids including tZR, IPA, and DCS was significantly elevated in *PtHMGR-OEs*, indicating that *PtHMGR* overexpression directly influences the biosynthesis of MVA-related isoprenoids. Therefore, the *HMGR* gene directly affects MVA-derived isoprenoids and indirectly affects the content of MEP-derived isoprenoids by changing the expression levels of MEP-related genes.

### 4.5. Higher levels of MEP- and MVA-derived products in PtDXR-OE seedlings

DXR is the rate-limiting enzyme in the MEP pathway and an essential regulatory step in the cytoplasmic metabolism of isoprenoid compounds (Takahashi et al., 1998). Mahmoud and Croteau (2001) revealed that overexpression of *DXR* in *Mentha piperita* promoted the synthesis of monoterpenes in the oil glands and increased the production of essential oil yield by 50%. However, the up-regulation of *DXR* expression did not lead to change in the complex oil composition significantly. Hasunuma et al. (2008) exhibited that overexpression of *Synechocystis sp*. strain PCC6803 *DXR* in tobacco resulted in increased levels of β-carotene, chlorophyll, antheraxanthin, and Lutein. Xing et al. (2010) showed that the *A*. *thaliana dxr* mutants caused to lack of GAs, ABA, and photosynthetic pigments (REF57). These mutants showed pale sepals and yellow inflorescences (Xing et al., 2010). In our study, the relatively higher abundance of GA3 and carotenoids in *PtDXR-OE* poplar seedlings indicated an effect of *DXR* overexpression. Combined with the result described above of increased *DXS*, *HDS*, *HDR*, *MCT*, *CMK*, *FPS*, *GPS*, and *GPPS* expression levels, we postulate that overexpression of *DXR* not only affects the expression levels of MEP-related genes but also changes the field of GA3, and carotenoids.

### 4.6. Communications exist between MVA- and MEP-pathways excess of IPP and DMAPP

Although the substrates of MVA- and MEP pathways differ, there are common intermediates like IPP and DMAPP (Figure 6). Blocking only the MVA or the MEP pathway, respectively, does not entirely prevent the biosynthesis of terpenes in the cytoplasm or plastids, indicating that some MVA and MEP pathways products can be transported and/or move between cell compartments (Aharoni et al., 2003; Aharoni et al., 2004; Gutensohn et al., 2013). For example, it has been shown that the transferring IPP from the chloroplast to cytoplasm observed through 13C labeling, indicating that plentiful IPP is available for use in the MVA-pathway to produce terpenoids (Ma et al., 2017). In addition, segregation between the MVA- and MEP pathways is limited and might exchange some metabolites over the plastid membrane (Laule, 2003). Kim et al. (2016b) used clustered, regularly interspaced short palindromic repeats (CRISPR) technology to reconstruct the lycopene synthesis pathway and control the flow of carbon in the MEP-and MVA-pathways. The results showed that the expression of MVA-related genes was reduced by 81.6%, but the lycopene yield was significantly increased. By analyzing gene expression levels and metabolic outcome in *PtHMGR*-and *PtDXR-OEs*, we discovered that the correlation might exist between MVA- and MEP-related genes with MVA- and MEP-derived products, which are not restricted to crosstalk between IPP and DMAPP (Figure 6).

**Figure 6.**
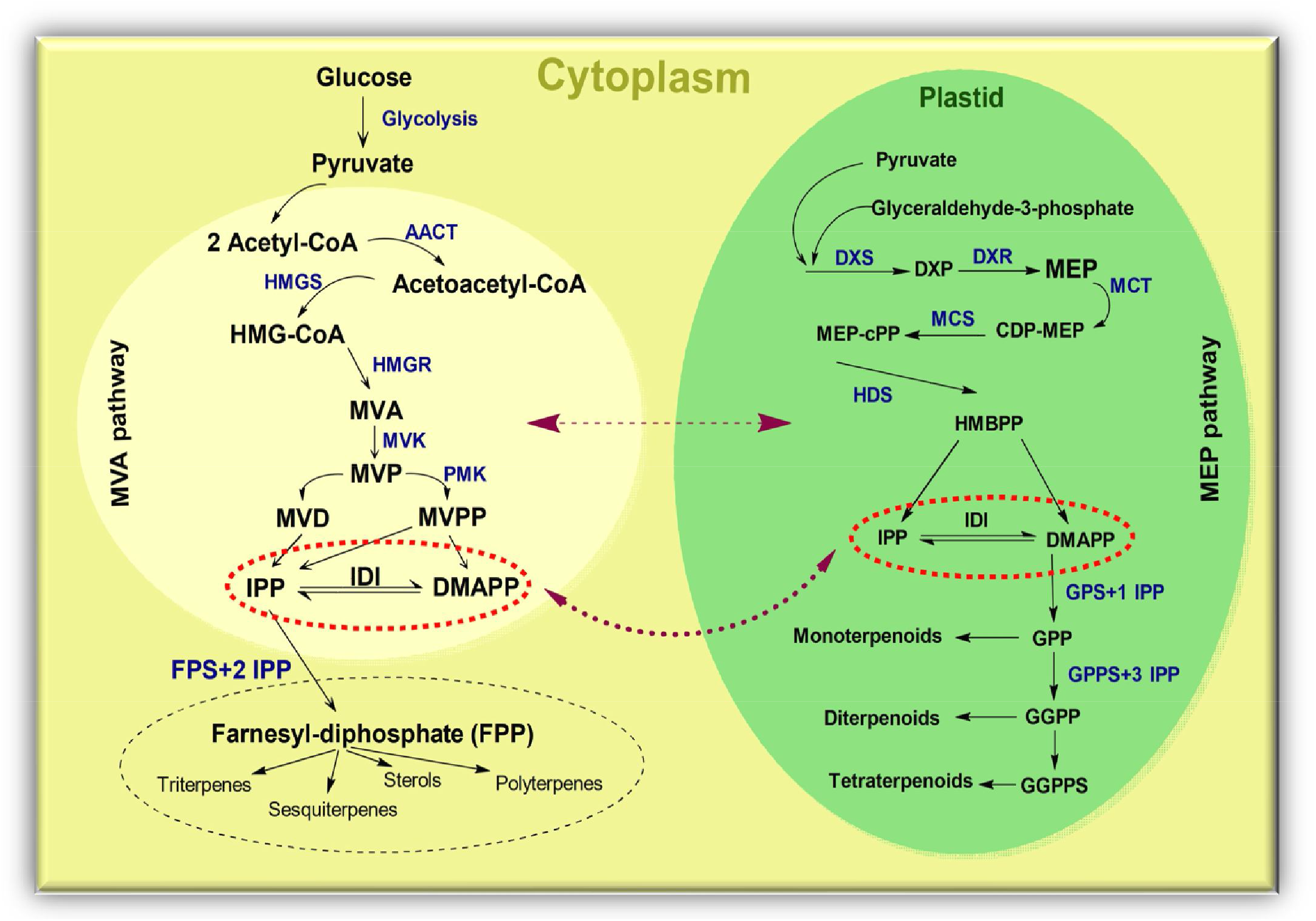
Communications exist between MVA- and MEP-pathways excess of IPP and DMAPP. The IPP and DMAPP are considered the common precursors of the MEP- and MVA pathways between cytoplasm and plastid. In addition, the putative communication generated between MVA- and MEP-related genes and MVA- and MEP-derived products. MVA: mevalonic acid, MEP: methylerythritol phosphate, IPP: isopentenyl diphosphate, DMAPP: dimethylallyl diphosphate, AACT: acetoacetyl CoA thiolase, HMGS: 3-hydroxy-3-methylglutaryl-CoA synthase, HMG-CoA: 3-hydroxy-3-methylglutary-CoA, HMGR: 3-hydroxy-3-methylglutaryl-CoA reductase, MVK: mevalonate kinase, MVD: mevalonate5-diphosphate decarboxylase, IPP: isopentenyl diphosphate, IDI: IPP isomerase, GPP: geranyldiphosphate, FPP: famesyldiphosphate, GPS: geranyl phosphate synthase, FPS: farnesyl-diphosphate synthase, GPPS: geranyl diphosphate synthase, GGPPS: geranyl geranyl diphosphate synthase, DXS: 1-deoxy-D-xylulose5-phosphate synthase, DXP: 1-deoxy-D-xylulose5-phosphate, DXR: 1-deoxy-D-xylulose5-phosphate reductoisomerase, HDS: 1-hydroxy-2-methyl-2-(E)-butenyl4-diphosphate synthase, HDR: 1-hydroxy-2-methyl-2-(E)-butenyl4-diphosphate reductase, MCT: MEP cytidylyltransferase, CMK: 4-diphosphocytidyl-2-C-methyl-D-erythritol kinase.

On the one hand, overexpression of *PtDXR* affected the transcript levels of MEP-related genes and the contents of MEP-derived isoprenoids, including GAs and carotenoids. The diminished accumulation of MVA-related gene products causes a reduction in the yields of MVA-derived isoprenoids (including CS) but leads to increasing tZR, IPA, and DCS contents. We hypothesize that IPP and DMAPP produced by the MEP pathway could enter the cytoplasm to compensate for the lack of IPP and DMAPP, and the IPP and DMAPP as the precursors of the MVA pathway are used to guide the synthesis of MVA-derived products. On the other hand, *PtHMGR-OEs* exhibited higher transcript levels of *AACT*, *MVK*, and *MVD* and higher *DXS*, *DXR*, *HDS*, and *HDR* than NT poplars, resulting in effect both MEP- and MVA-related gene expressions. We successfully demonstrated that manipulation of *HMGR* in the poplar MVA pathway results in dramatically enhanced yields of GAs and carotenoids. This result illustrates that cytosolic *HMGR* overexpression expanded plastidial GPP- and GGPP-derived products, such as carotenoids. Therefore, this study provides hints that communications between the MVA-and MEP pathways increased the expression levels of *GPS* and *GPPS* in *PtHMGR-OEs*, and elevated the contents of GA3 and carotenoids. Moreover, changes in MEP- and MVA-related gene expressions affect MVA- and MEP-derived isoprenoids.

In conclusion, overexpression of *PtHMGR* in poplars caused the accumulation of MVA-derived isoprenoids and MEP-derived substances. The advanced insights into the regulation of MVA- and MEP pathways in poplar add to the knowledge about these pathways in Arabidopsis, tomato, and rice. In *PtHMGR-OE* poplars, most MEP- and MVA-related genes associated with the biosynthesis of isoprenoid precursors were upregulated. In *PtDXR-OE* poplars, elevated contents of GAs, carotenoids, and GRs were attributed to increased expression of MEP-related genes as well as plastidial *GPP* and *GGPP*. Together, these results show that manipulating *PtDXR and PtHMGR* is a novel strategy to influence poplar isoprenoids.

### 4.7. Communications between MVA- and MEP pathways affected by PtHMGR- and PtDXR-OEs influence the plant growth and developments

It has been shown that Abscisic acid (ABA) and GA3 perform essential functions in cell division, shoot growth, and flower induction (Xing et al., 2016). It has also been demonstrated that the cytokinin tZR, a variety of phytohormones, perform a crucial function as root to shoot signals, directing numerous developmental and growth processes in shoots (Abul et al., 2010; Sakakibara, 2006). Regarding these findings, we showed how the communications between MVA- and MEP pathways and their changes affected by some stimulators (*HMGR*-and *DXR-OEs*) influenced plant growth, especially in stem length. Finally, We figured out that the gibberellic acid and cytokinin may be more effective in plant growth than inhibiting by ABA, causing higher *PtDXR-OEs* than *PtHMGR-OEs* compared with NT poplars.

## Author contributions

A.M. and H.W. conceived, planned, and coordinated the project, performed data analysis, wrote the draft, and finalized the manuscript. B.P. validated and contributed to data analysis and curation, revised and finalized the manuscript. T.J., W.S., and D.L. reviewed and edited the manuscript. L.Y. and Q.Z. coordinated, contributed to data curation, finalized and funded this research. A.M., H.W., and B.P. contributed equally as the first author.

## Acknowledgments

This work was supported by the National Key Program on Transgenic Research (2018ZX08020002), the National Natural Science Foundation of China (No. 31971682).

## Supplementary figures and table

**Supplementary Figure 1 | Amino acid sequences alignment of PtHMGR protein and other known HMGR proteins.** *A. thaliana* (NP_177775.2), *G. hirsutum* (XP_016691783.1), *M. domestica* (XP_008348952.1), *M. esculenta* (XP_021608133.1), *P. persica* (XM_020569919.1), *O. sativa* (XM_015768351.2), *T. cacao* (XM_007043046.2), *Z. mays* (PWZ28886.1). The HMG-CoA and NADPH binding domains are indicated in red rectangular.

**Supplementary Figure 2 | Construction of a phylogenetic tree based on the HMGR sequences of various species.** Accession numbers of the HMGR obtained from Phytozome are as follows: *A. thaliana* (AT1G76490 and AT2G17370), *P. trichocarpa* (Potri.011G145000, Potri.005G257000, Potri.004G208500, Potri.001G457000, Potri.009G169900 and Potri.002G004000), *Gossypium raimondii* (Gorai.008G013000, Gorai.002G146000, Gorai.002G014700, Gorai.005G215800, Gorai.012G138100, Gorai.005G215500, Gorai.005G215600 and Gorai.005G215700), *Malus domestica* (MDP0000157996, MDP0000268909, MDP0000372490, MDP0000251253 and MDP0000312032), *Manihot esculenta* (Manes.15G114100, Manes.01G157500, Manes.03G096600, Manes.02G116900 and Manes.05G128600), *Oryza sativa* (LOC_Os09g31970, LOC_Os08g40180 and LOC_Os02g48330), *Prunus persica* (Prupe.7G187000, Prupe.7G187500 and Prupe.8G182300), *Theobroma cacao (*Thecc1EG000025, Thecc1EG007601 and Thecc1EG034814), and *Zea mays* (GRMZM2G393337, GRMZM2G058095, GRMZM2G136465, GRMZM2G001645 and GRMZM2G043503).

**Supplementary Figure 3 | Molecular identification of *PtHMGR-OEs*.** (**A**) PCR identification of *PtHMGR* in *PtHMGR-OEs* and NT poplars. Lane M: 15K molecular mass marker (TransGen, China); lane 1: genome DNA from WT as negative control; lanes 2–9: genome DNA from *PtHMGR-OE* lines (**B**) qRT-PCR identification of the transcript levels of *PtHMGR* in *PtHMGR-OEs* and NT poplars. Three independent experiments were performed; Stars reveal significant differences, * P < 0.05, ** P < 0.01, *** P < 0.001.

**Supplementary Figure 4 | MEP- and MVA-related genes analyses in overexpressed *PtHMG R* - and *PtDXR-OEs* poplars. a**, MVA-relate genes *AACT*, *HMGS*, *MVK*, *MVD*, and *FPS* affected b y *PtHMGR* overexpressing. **b**, MVA-related genes *AACT*, *HMGS*, *HMGR*, *MVK*, *MVD*, and *FPS* affected by *PtDXR* overexpressing. **c**, MEP-related genes *DXS*, *MCT*, *CMK*, *HDS*, *HDR*, *IDI*, *GPS, G PPS, and DXR affected by PtHMGR* overexpressing. **d**, MEP-related genes *DXS*, *DXR*, *MCT*, *CM K*, *HDS*, *HDR*, *IDI*, *GPS*, and *GPPS* affected by *PtDXR* overexpressing. *PtActin* was used as an in ternal reference in all repeats; * P < 0.05, ** P < 0.01, ***P < 0.001, ****P < 0.0001; Three in dependent replications were performed in this experiment.

**Supplementary Figure 5 |** HPLC chromatograms of analyzing the contents of **(A)** β-carotene, **(B)** lycopene, and **(C)** lutein in NT poplars and *PtHMGR-OEs*.

**Supplementary Figure 6 |** HPLC chromatograms of analyzing the contents of **(A)** β-carotene, **(B)** lycopene, and **(C)** lutein in NT poplars and *PtDXR-OEs*.

**Supplementary Figure 7 | Chromatogram analyses of GA3 standards via HPLC-MS/MS.** The chromatogram of standard GA3 at (**A**) 0.1, (**B**) 0.2, (**C**) 0.5, (**D**) 2, (**E**) 5, (**F**) 20, (**G**) 50, and (**H**) 200 ng/mL concentrations. (**I**) Equations for the GA3 standard curves.

**Supplementary Figure 8 | Chromatogram analyses of tZR standards via HPLC-MS/MS.** The chromatogram of standard tZR at (**A**) 0.1, (**B**) 0.2, (**C**) 0.5, (**D**) 2, (**E**) 5, (**F**) 20, (**G**) 50, and (**H**) 200 ng/mL concentrations. (**I**) Equations for the tZR standard curves.

**Supplementary Figure 9 | Chromatogram analyses of IPA standards via HPLC-MS/MS.** The chromatogram of standard IPA at (**A**) 0.2, (**B**) 0.5, (**C**) 2, (**D**) 5, (**E**) 20, (**F**) 50, and (**G**) 200 ng/mL concentrations. (**H**) Equations for the IPA standard curves.

**Supplementary Figure 10 | Chromatogram analyses of DCS standards via HPLC-MS/MS.** The chromatogram of standard DCS at (**A**) 0.5, (**B**) 2, (**C**) 10, (**D**) 20, and (**E**) 50 ng/mL concentrations. (**F**) Equations for the DCS standard curves.

**Supplementary Figure 11 | Chromatogram analyses of CS standards via HPLC-MS/MS.** The chromatogram of standard CS at (**A**) 0.5, (**B**) 5, (**C**) 10, (**D**) 20, and (**E**) 50 ng/mL concentrations. (**F**) Equations for the CS standard curves.

**Supplementary Table 1 |** Primers were used in this study.

**Supplementary Table 2 |** Table of data analyses used in phenotypic changes evaluation. **a**, Stem diameter data analyses. **b**, Stem length data analyses.

